# Evaluation of single-cell RNA profiling technologies using FFPE, fresh, and frozen ccRCC tumor specimens

**DOI:** 10.1101/2025.11.25.690485

**Authors:** Sherin Xirenayi, Sabrina Y. Camp, Amanda E. Garza, Yashika Rustagi, Erica Pimenta, Erin Shannon, Anwesha Nag, Adam Betherman, Anthony Anselmo, Rachel Trowbridge, Aaron R. Thorner, Jihye Park, Kevin Bi, Eliezer M. Van Allen

## Abstract

The application of single-cell transcriptomic approaches has deepened our understanding of tumor heterogeneity, immune dynamics, and molecular programs underlying therapy response. The recent development of fixation-compatible single-cell platforms, such as 10x Genomics Flex, offers the opportunity to profile archived formalin-fixed, paraffin-embedded (FFPE) specimens, expanding access to clinically valuable samples. However, most benchmarking studies of recent single-cell RNA sequencing (scRNA-seq) technologies have relied on peripheral blood mononuclear cells, limiting their relevance to human tissues. Here, we compared three 10x single-cell RNA profiling methods, fresh tumor scRNA-seq, flash-frozen single-nucleus Multiome, and FFPE single-nucleus Flex (snFlex), using clear cell renal cell carcinoma (ccRCC) biopsies. Across methods, we observed broadly consistent cell type-specific transcriptional profiles among major ccRCC cell populations. Despite lower gene and UMI counts, snFlex reliably identified fine-grained states within CD8^+^ T cells, tumor-associated macrophages, and tumor compartments, comparable to those detected by scRNA-seq. Together, these findings highlight the distinct advantages of each technology depending on sample preservation type and study design, providing practical guidance for single-cell RNA profiling technology selection in translational studies using human tumor biopsies.

## INTRODUCTION

Recent advances in clinically integrated single-cell transcriptomic profiling have transformed translational cancer research by revealing cellular heterogeneity within tumors, identifying immune subtypes linked to clinical outcomes, and uncovering transcriptional tumor states associated with therapy response^1–4^. For example, Bi et al. profiled 8 renal cell carcinoma lesions and identified key immune axes, including CXCL10-high macrophages and 4-1BB-high CD8^+^ T cells, that were associated with different immunotherapy outcomes^1^. Similarly, He et al. analyzed 14 metastatic castration-resistant prostate cancer biopsies and identified distinct transcriptional states, including cancer intrinsic epithelial-mesenchymal transition programs and clonally expanded dysfunctional CD8^+^ T cells associated with treatment resistance, providing a foundation for developing complementary therapeutic strategies^2^.

However, the application of single-cell RNA sequencing (scRNA-seq) to primary human tumors remains challenging because this strategy requires freshly dissociated specimens, which are difficult to obtain and must be processed immediately after acquisition. Moreover, samples collected at different times or under variable conditions often introduce technical heterogeneity that can confound biological interpretation. These limitations make it labor-intensive to implement scRNA-seq in clinical settings and introduce logistical barriers for large-scale or longitudinal studies.

Single-nucleus RNA sequencing (snRNA-seq) from flash-frozen specimens offers an alternative that eliminates the need for fresh tissue; however, it typically yields reduced immune cell recovery^5,6^. This poses a major limitation, as immune cell abundance and transcriptional states have been associated with prognosis and therapeutic outcomes in several cancer types. Recently developed formalin-fixed, paraffin-embedded (FFPE)-compatible single-cell RNA sequencing platforms, such as the 10x Genomics Flex, provide an opportunity to profile transcriptomes from archived clinical samples and potentially overcome these challenges; however, the relative capacity of recovering relevant biological information important for tumor-based inquiries is uncertain, as most benchmarking studies have relied on peripheral blood mononuclear cells.

To address this gap, we performed a comparative analysis of three 10x single-cell RNA profiling modalities—fresh-tumor scRNA-seq, flash-frozen single-nucleus Multiome (RNA modality), and FFPE single-nucleus Flex (snFlex)—using clear cell renal cell carcinoma (ccRCC) biopsies, evaluating library quality and complexity, marker gene expression, and cell type recovery. ccRCC is the most common subtype of renal cell carcinoma and is highly immune-infiltrated^7,8^, and recent studies have highlighted immune-mediated mechanisms of therapy resistance^1,9–11^, making immune cells one of the key components to study in this aggressive cancer and a useful histology to assess single-cell profiling modalities and considerations across cancers. Across methods, we observed broadly consistent cell type-specific transcriptional profiles among major cell populations. Despite lower gene and UMI counts, snFlex reliably identified fine-grained states within CD8^+^ T cells, tumor-associated macrophages, and tumor compartments, comparable to those detected by scRNA-seq. Together, these findings highlight the distinct advantages of each technology depending on sample preservation type and study design, providing practical guidance for single-cell RNA profiling technology selection in translational studies using human tumor biopsies.

## RESULTS

### snFlex, scRNA, and snMultiome capture broadly similar cell type-specific transcriptional profiles across major cell types

To evaluate whether FFPE-based single-cell RNA profiling of ccRCC tumor specimens can recover meaningful biological insights, we compared data generated using three 10x Genomics technologies that differ in tissue preservation and sequencing chemistry: (1) FFPE-derived samples processed with 10x Flex technology, which uses single-nucleus isolation and probe-based chemistry (referred to as snFlex); (2) freshly dissociated tumors using conventional single-cell 3′ capture chemistry (scRNA); (3) flash-frozen tissues profiled with 10x Multiome using single-nucleus isolation and 3′ capture chemistry, from which RNA expression data were used in this study (referred to as snMultiome). We analyzed 12 ccRCC samples that had been previously processed and sequenced by our group^1,12^ and generated an additional 4 ccRCC datasets using snFlex in collaboration with the DFCI Center for Cancer Genomics (CCG). Among these, one patient (patient 876) had primary tumor biopsies profiled using all three methods. We focused on this patient for direct method comparison, including cell type-specific marker gene expression, cell type recovery, and granular phenotyping of immune and tumor populations, and incorporated the full cohort to validate our findings (Figure 1A).

**Figure 1.**
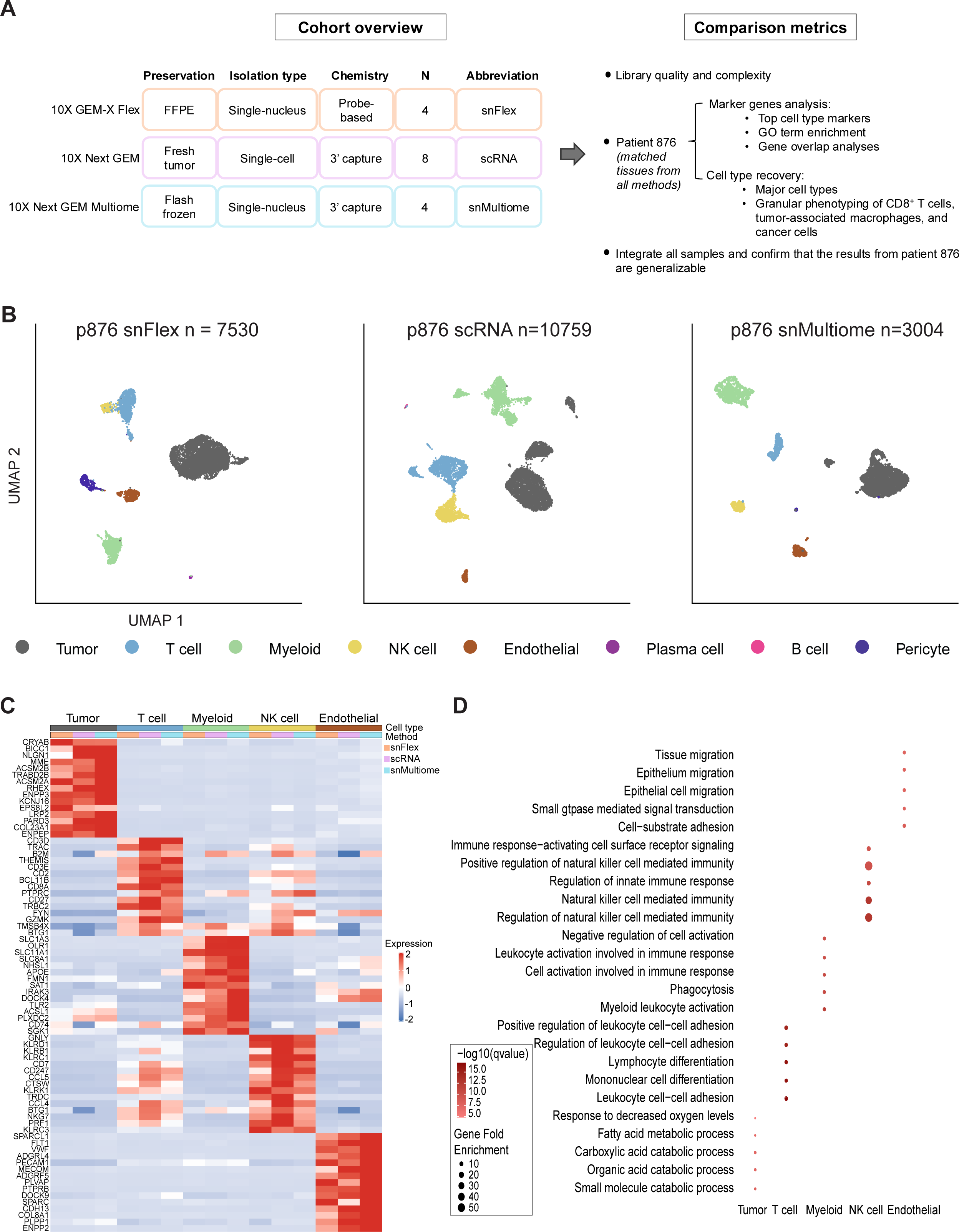
snFlex, scRNA, and snMultiome capture broadly similar cell type-specific transcriptional profiles across major cell types. A. Cohort overview and study schematics. B. Uniform manifold approximation and projection (UMAP) of tumor, stromal, and immune cells captured in patient 876 (p876) across methods, colored by broad cell type. C. Heatmap showing average expression of the top 15 marker genes shared across methods for each common cell type in patient 876. Marker genes were identified using one-versus-all logistic regression differential expression (DE) tests (Bonferroni-adjusted p < 0.05) performed within each shared cell type for each sequencing method while accounting for sequencing depth. For each cell type, genes identified as markers in all methods were defined as shared markers, ranked by adjusted p-value and log fold change, and the top 15 were plotted. Columns are grouped by cell type and annotated by both cell type and sequencing method. D. Dot plot of pathways enriched among shared cell type marker genes across sequencing methods in patient 876, identified with clusterProfiler based on Gene Ontology (GO) Biological Process terms (Benjamini–Hochberg FDR < 0.05). Dot size indicates gene fold enrichment (GeneFoldEnrich), dot color represents statistical significance (-log10 q value). The top five enriched pathways for each cell type are displayed.

After quality control and cell type annotation, patient 876’s samples yielded 7,530, 10,759, and 3,004 high-quality cells from snFlex, scRNA, and snMultiome, respectively. Each method robustly captured the major cellular compartments of ccRCC, including tumor, immune (T, myeloid, NK), and stromal (endothelial) populations. Some populations showed method-specific recovery. For instance, plasma B cells characterized by *JCHAIN* and *IGKC* were observed in snFlex and scRNA, but not in snMultiome. B cells, marked by *CD79A* and *MS4A1*, were unique to scRNA, and pericytes were detected in snFlex and snMultiome, but not in scRNA (Figure 1B, S1A; Methods). Major ccRCC cell types were also consistently detected across all samples in the integrated dataset (Figure S2A, B)

To determine whether the three methods yielded broadly similar cell type-specific transcriptional markers, we compared marker genes for cell types captured across all methods. In single-patient analyses, to enable a direct method-level comparison while accounting for differences in total cell numbers, we proportionally downsampled snFlex and scRNA to match the total number of cells in snMultiome (2,972) while maintaining their original cell type proportions (Methods). Within the downsampled snFlex and scRNA datasets and in snMultiome, we identified differentially expressed genes (DEGs) for each shared cell type. The top 15 shared cell type-specific genes generally aligned with well-established, cell type-specific markers (Figure 1C; Table S1D). For each method, we also examined its top 15 markers and found that they aligned with expected lineage signatures and were robustly detected across platforms, except for long non-coding RNAs (lncRNAs) and Human Leukocyte Antigen (HLA) genes, which were not captured in snFlex, due to probe design constraints (Figure S1B; Tables S1E-G).

We next asked whether this trend held across the full cohort. For each method, we identified DEGs for every cell type without downsampling to capture broader transcriptional patterns across samples, recognizing that these comparisons inherently reflect both biological and technical heterogeneity. To mitigate these effects, we applied batch correction using Harmony and performed logistic regression differential expression analysis with patient identity as a covariate (Methods). The top 15 cell type-specific genes shared across methods again corresponded to canonical markers (Figure S2C; Table S2B).

Finally, we performed gene set enrichment analysis (GSEA) using Gene Ontology (GO) biological process terms to evaluate whether the shared markers reflected expected cell type functional programs across methods. Each cell type’s marker set was enriched for its expected functional programs in both the single-patient and integrated analyses (Figure 1D, S2D; Tables S1H, S2F). For example, tumor cells showed enrichment in metabolic processes, T cells in lymphocyte differentiation, and myeloid cells in myeloid leukocyte activation and phagocytosis pathways. Overall, despite differences in tissue preservation and chemistry, all three technologies captured the major ccRCC cell types and consistently reflected their canonical biological functions.

### Library complexity, cell type composition, and marker gene expression vary across methods

When comparing library complexity and quality, we observed that snFlex captured fewer genes and unique molecular identifiers (UMIs) than scRNA or snMultiome, whereas scRNA exhibited the highest percentage of mitochondrial reads per cell in both the single-patient and integrated analyses (Figure 2A, S3A). We next examined the proportion of each cell type identified in patient 876 and across the full cohort and observed a consistent trend. Tumor and stromal cells were enriched in snFlex, immune cells were more abundant in scRNA, and snMultiome showed a more balanced recovery across cell types (Figure 2B, S3B).

**Figure 2.**
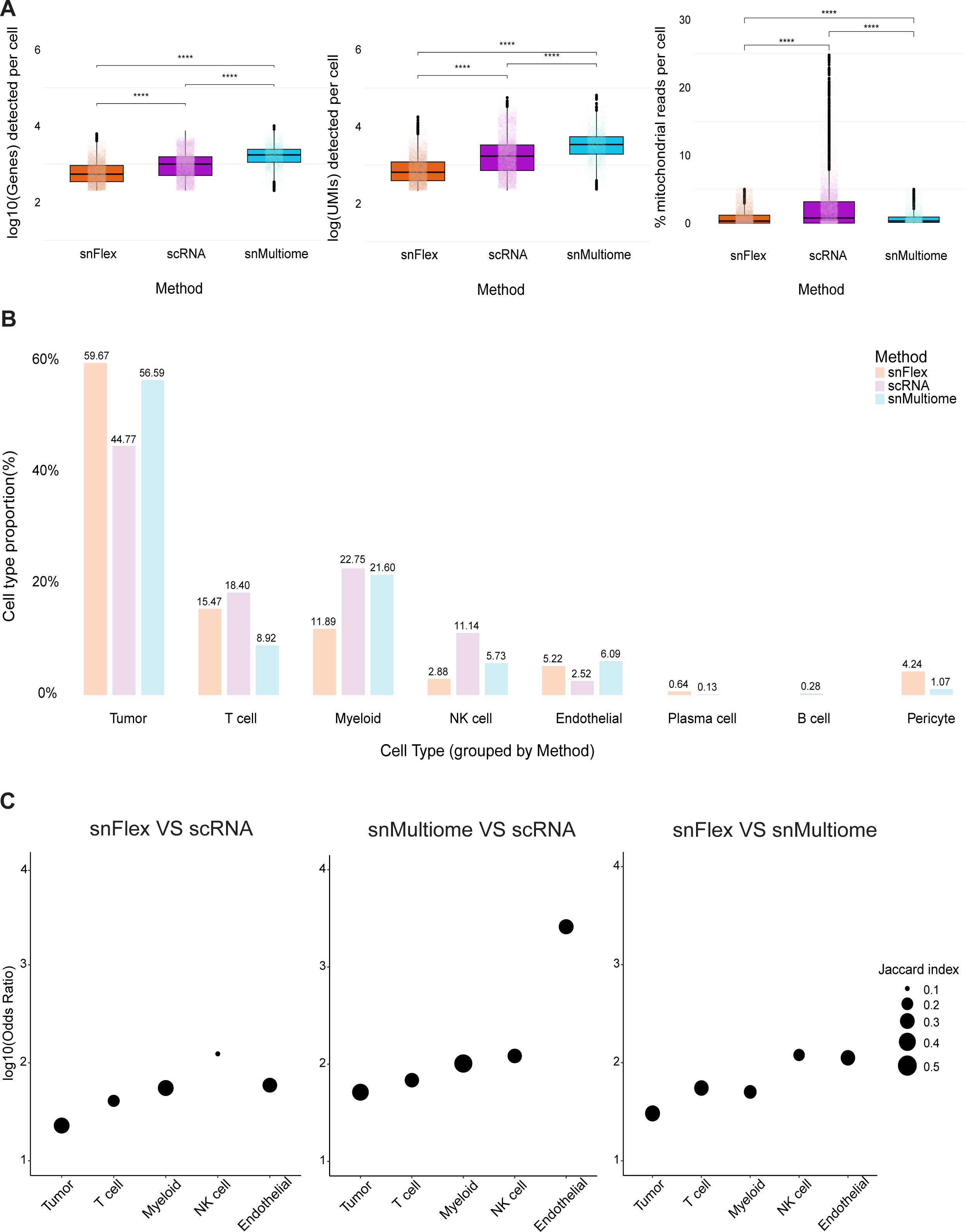
Library complexity, cell type composition, and marker gene expression vary across methods. A. Box plots showing the number of detected genes, unique molecular identifiers (UMIs), and the percentage of mitochondrial reads per cell across sequencing methods for patient 876. Gene and UMI counts are displayed on a log_10_ scale. Statistical significance was assessed using a two-sided Wilcoxon rank-sum test. B. Bar plot showing the proportion of each cell type identified in patient 876 across sequencing methods, with proportions calculated within each method and grouped by cell type. C. Dot plots showing gene overlap analyses of cell type markers across sequencing methods, evaluated using Fisher’s exact test and Jaccard similarity. The y-axis shows the log₁₀of odds ratios from Fisher’s exact test, representing enrichment strength. All overlaps were statistically significant (p < 0.0001). Dot size indicates the Jaccard index, and all panels are plotted on consistent scales.

To quantitatively assess similarities and differences in marker genes across shared cell types, we performed pairwise per-cell type gene overlap analyses between methods using Fisher’s exact test and Jaccard similarity, based on cell type-specific markers identified within each method (Methods). In patient 876, all pairwise comparisons were highly significant (p < 0.0001; Fisher’s exact test), with tumor markers showing the lowest odds ratios (OR), T and myeloid cells at intermediate levels, and NK and endothelial cells with higher OR. The Jaccard similarities were generally consistent, with most cell types having a moderate overlap (0.2 – 0.4), except NK cells between snFlex and scRNA, which showed a lower similarity of 0.1 (Figure 2C). Across the full cohort, a similar overall pattern was observed. All pairwise comparisons were highly significant (p < 0.0001; Fisher’s exact test), with tumor, T, and myeloid populations displaying lower OR and stromal populations exhibiting stronger concordance across methods. The Jaccard overlaps consistently indicated moderate to high similarities (0.2 – 0.6) across all comparisons (Figure S3C). Overall, these analyses revealed method-specific differences in library complexity and cell type composition, but also demonstrated a relatively consistent agreement of cell type markers across methods.

### snFlex enables high-resolution phenotyping of immune and tumor subsets

Since one of the key advantages of scRNA is its ability to capture granular and rare cell states, we then asked whether snFlex could achieve comparable resolution in identifying transcriptionally distinct populations for cancer applications, taking scRNA as the ground truth. We focused on CD8^+^ T cells, tumor-associated macrophages (TAMs), and tumor cells. Each granular phenotype was defined as described in the Methods section (in patient 876, there were not enough T cells in snMultiome to perform detailed phenotyping, so we compared snFlex and scRNA in that setting).

In the single-patient analysis of CD8^+^ T cells, both methods revealed three distinct CD8^+^ T cell states previously described in ccRCC^1^. A 4-1BB-high population characterized by elevated co-stimulatory, effector, and inhibitory receptor gene expression, a 4-1BB-low cluster enriched for memory and longevity-associated genes, and a cycling subset representing proliferative CD8^+^ T cells (Figure 3A; Tables S3A-C). To further define the 4-1BB-high and 4-1BB-low CD8^+^ T cell states, we performed GSEA analysis using T cell signatures from Oliveira et al^13^ (Methods; Tables S3D, E). As expected, the 4-1BB-high clusters in both snFlex and scRNA were enriched for terminal exhaustion signatures, with comparable enrichment scores. The overlapping leading-edge genes, such as *ENTPD1*, *TNFRSF18*, and *LAYN*, are well-established markers of terminally exhausted CD8^+^ T cells^1,13–16^. Similarly, the 4-1BB-low clusters in both methods were enriched for effector memory-associated signatures, with overlapping leading-edge genes corresponding to canonical T cell memory markers such as *SELL*, *IL7R*, and *CCR7*^1,13,16,17^(Figure 3B). Together, these analyses show that snFlex recapitulates the biologically relevant phenotypic states observed by scRNA.

**Figure 3.**
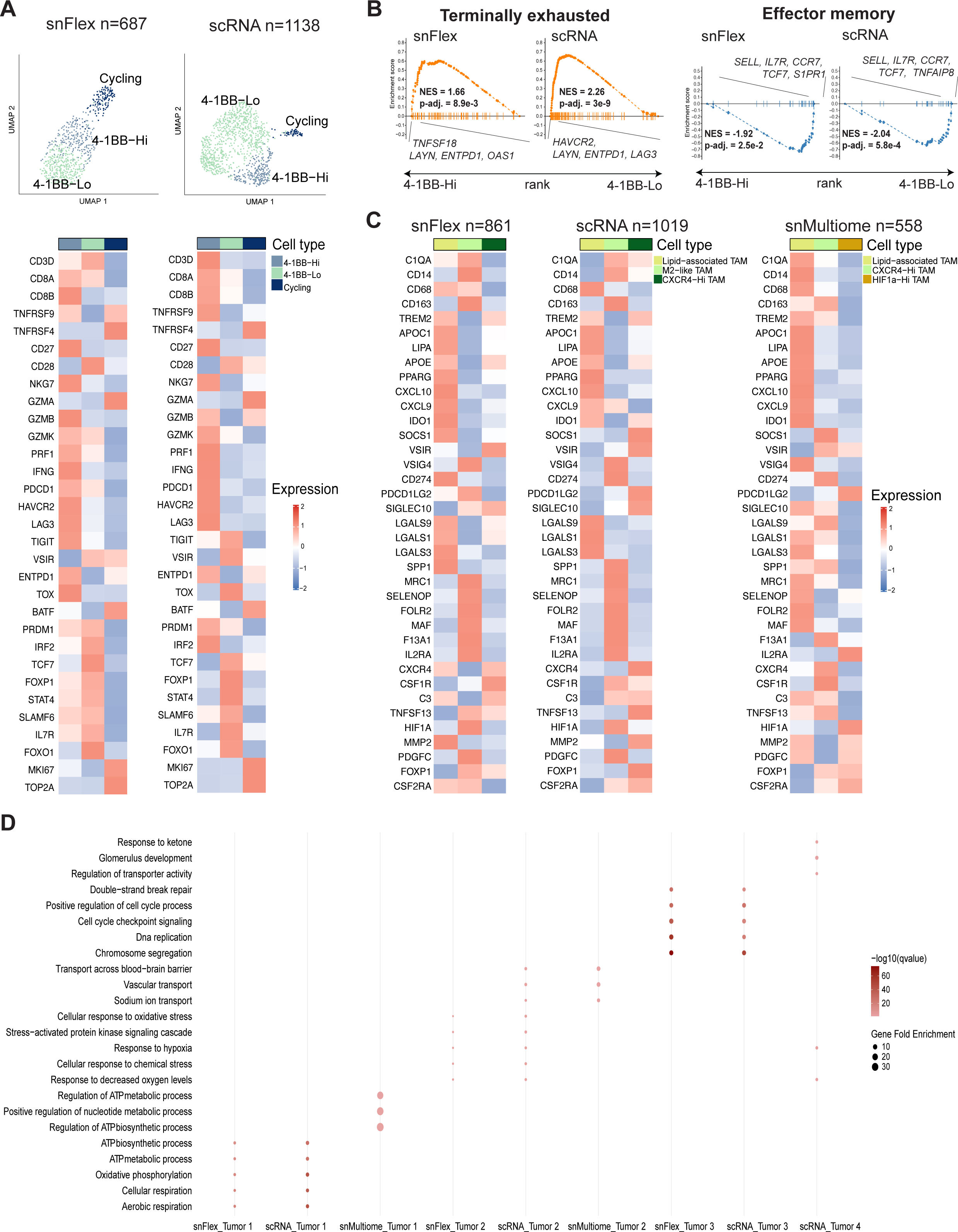
snFlex enables high-resolution phenotyping of immune and tumor subsets. A. UMAP visualization of granular CD8^+^ T cell states captured by snFlex and scRNA in patient 876 (top), followed by heatmaps showing scaled expression of the same set of cell state-defining genes (bottom). Marker genes were identified by Multinomial-based Alignment of Single-cell Transcriptomics (MAST) model and two-sided Wilcoxon rank-sum tests (Bonferroni-adjusted p < 0.05). B. Gene set enrichment analysis (GSEA) of terminally exhausted (left) and effector memory (right) CD8⁺ T cell signatures from Oliveira et al., 2021 (Nature), comparing 4-1BB-Hi and 4-1BB-Lo clusters across snFlex and scRNA. Leading-edge genes, normalized enrichment score (NET), and p-adj value are shown on each plot. C. Heatmaps showing scaled expression of the same set of genes defining granular tumor-associated macrophage (TAM) phenotypes across methods in patient 876, identified by MAST model and two-sided Wilcoxon rank-sum tests (Bonferroni-adjusted p < 0.05). D. Dot plot showing pathways enriched from granular tumor marker genes across sequencing methods in patient 876. Pathway enrichment was performed using clusterProfiler based on GO Biological Process terms (Benjamini–Hochberg FDR < 0.05). Marker genes were identified using MAST and two-sided Wilcoxon rank-sum test (Bonferroni-adjusted p < 0.05). Dot size represents fold enrichment, and dot color indicates statistical significance (–log₁₀ q-value). Displayed pathways were manually selected from significant GO Biological Process terms.

The integrated analysis across all samples revealed six CD8^+^ T cell states, including 4-1BB-high, interferon-stimulated gene (ISG)-high, cycling, early activated-like, scRNA-dominant, and NK-like populations. Among these, the 4-1BB-high, ISG-high, cycling, and early activated-like CD8^+^ T cells were consistently represented across all three methods (Figure S4A). To visualize these states across methods, we selected representative functional genes for each state from the DEGs defining shared CD8^+^ T cell clusters in the integrated dataset and summarized their average expression in a heatmap^1,9,16,18^ (Figure S4B; Methods; Table S4A). This analysis revealed that all three technologies reliably captured shared CD8⁺ T cell subtypes.

In the single-patient analysis of TAMs, both snFlex and scRNA identified three subtypes. A population of lipid-associated macrophages expressing elevated lipid metabolism and immune-suppressive markers^19^, an M2-like subtype expressing genes associated with the M2 phenotype^1,20^, and a CXCR4-high cluster enriched for migration and survival genes^19^. snMultiome captured the lipid-associated and CXCR4-high subsets but not a distinct M2 cluster, though the associated genes were expressed. Instead, this modality revealed a HIFα-high population enriched for hypoxia and angiogenesis pathways^21^ (Figure 3C; Table S5). Overall, the macrophage subtypes were largely consistent in patient 876, with the highest concordance observed between snFlex and scRNA. Across the full cohort, we identified six distinct TAM populations, including CXCL10-high, lipid-associated, M2-like, cycling, lncRNA-high, and lipid-associated SPP1-high. Only the lipid-associated SPP1-high subtype was uniquely captured in scRNA, whereas the others were shared across methods (Figure S5A). Using the same approach as in the CD8^+^ T cell analysis, we compared the expression of functional TAM-defining genes and found that CXCL10-high and lipid-associated TAMs displayed similar gene expression profiles across methods^9,19,20^. The M2-like population was more prominently represented in snFlex, and lncRNA-high subset was enriched in snMultiome (Figure S5B; Table S4B).

All three methods captured broadly convergent tumor programs in patient 876. For example, we observed a tumor state enriched for metabolic programs (tumor 1) across all three methods, a hypoxia enriched tumor state (tumor 2) in just snFlex and scRNA. scRNA also showed additional enrichment for genes encoding proximal tubule transporters, which was similarly observed in snMultiome. Both snFlex and scRNA contained a cycling tumor program, whereas scRNA uniquely captured a hypoxia-related cluster enriched for renal function pathways, including glomerular development and response to ketones (Figure 3D; Tables S6D-F). Across the full cohort, we identified four main tumor clusters (Figure S6A). A proximal tubule-like population defined by genes associated with renal proximal tubule function^22^, an oxidative metabolism-high subset enriched for oxidative metabolism pathways^1^, an interferon-stimulated cluster characterized by high ISG expression, and a cycling population representing proliferative tumor cells, along with an additional lncRNA-high cluster unique to scRNA (Figure S6B; Table S4C). Three of the four shared clusters exhibited similar expression of defining markers across methods. The interferon-stimulated tumor cluster in snFlex displayed higher expression of type I ISGs compared to the other methods, although all three platforms showed elevated expression of interferon-stimulated inhibitory or evasion markers^9,18^. Overall, each method supports a similar phenotypic characterization of ccRCC tumor clusters.

## DISCUSSION

Efforts to benchmark recent advances in commercially available single-cell RNA profiling technologies have largely relied on peripheral blood mononuclear cells^23,24^. While these studies offer well-controlled experimental systems suited for systematic method evaluation, their findings do not always translate directly to intact human tissues. In translational cancer research using human biopsies, experimental variability is common due to differences in sample handling, processing times, and data integration from multiple sources. Biological variability also arises because patient-derived specimens can originate from distinct treatment time points, anatomical sites, or clinical contexts, and are sometimes limited in number.

This study provides insight from two complementary perspectives. First, our cohort reflects these real-world complexities, which we addressed to the best of our ability by carefully evaluating both technical and biological variables and incorporating them as covariates in our analyses. Second, we include a unique case in which a single patient’s primary tumor was profiled using all three technologies, allowing a direct, within-patient comparison. Although samples were processed years apart and handled by different individuals, this design represents, to our knowledge, the first comparison of FFPE, fresh, and frozen single-cell profiling approaches performed on a single patient’s primary kidney tumor biopsies.

Across all three methods, we observed broadly consistent cell type-specific transcriptional signals among major ccRCC cell populations. Notably, snFlex, despite lower gene and UMI counts relative to scRNA-seq and snMultiome, reliably identified fine-grained states within CD8^+^ T cells, tumor-associated macrophages, and tumor compartments. However, given the limited sample size and inherent biological and technical variability of human tumors, the observed differences may be patient- or tumor type-specific.

Each single-cell sequencing technology offers distinct advantages depending on sample type and research goals. snFlex provides access to archived FFPE samples, and its fixation-based workflow enables multiplexed processing, improving efficiency and reducing batch effects. scRNA delivers the highest data quality for immune profiling, while snMultiome enables frozen-sample analysis with joint chromatin and transcriptome measurements. Limitations include snFlex’s probe-based design, which does not capture the full transcriptome; scRNA reliance on fresh tissue and potential induction of stress-related transcriptional programs during dissociation^25^; and reduced immune marker recovery in snMultiome. However, we cannot yet determine whether the hypoxia-associated transcriptional program observed in the scRNA dataset reflects biological or technical differences, as we did not analyze fresh scRNA datasets from other tumor types and project our hypoxia signature to evaluate whether similar patterns occur outside of ccRCC.

In conclusion, our study demonstrates that snFlex captures immune cell detail comparable to scRNA while offering practical advantages for studies where fresh tissue is unavailable. The choice of technology should ultimately be guided by research goals, tissue availability, and the biological questions being addressed.

## ACKNOWLEDGEMENTS

We thank the patients who participated in this study. This project was supported by NIH R01CA278980, R01CA279221, R50CA265182, and FUJIFILM Corporation.

## AUTHOR CONTRIBUTIONS

Conceptualization: SX, SYC, KB, EMVA; Data curation: SX, AEG, RT, JP; Formal analysis: SX, SYC; Funding acquisition: SX, JP, EMVA; Investigation: YR, AN, AB, AA, ART; Project administation: SX, AEG, ES, RT, JP; Resources: ART, EMVA; Supervision: ART, JP, KB, EMVA; Validation: SX; Visualization: SX; Writing -- original draft: SX, SYC, JP, KB, EMVA; Writing -- review & editing: AEG, YR, EP, ES, AN, AB, AA, RT, ART

## METHODS

### Sample collection, dissociation, library preparation, and sequencing for snFlex

#### Sample preparation and quality control

We sent four FFPE renal carcinoma tumor tissue specimens from patients 876, 909, 855, and 890 as scrolls to CCG. Total RNA was isolated from two 10-micron scrolls of each FFPE renal carcinoma tumor tissue sample (patient 876, patient 909, patient 855, and patient 890), using the RNeasy FFPE Kit (Catalogue no. 73504, Qiagen). RNA quality was assessed by calculating the DV-200 value using the RNA 6000 Pico Kit for the Agilent Bioanalyzer 2100 Systems. Samples with a DV-200 > 20% were included for single-cell isolation (patient 876: 20%, patient 855: 33%, patient 909: 30%; patient 890 failed with 1%).

#### Single-cell isolation

Three qualified samples were processed for single-cell isolation. Two 25-micron scrolls from patient 876 and patient 909, and four scrolls from patient 855 (split into two independent sub-samples, patient 855-a and patient 855-b) were deparaffinized twice with xylene and rehydrated sequentially with decreasing concentration of ethanol (100%, 70%, 50%) and sterile milli-Q water at RT. Finally, the cells were kept in sterile 1X Phosphate Buffer Saline (PBS) and incubated on ice.

Tissue dissociation was performed by incubating the scrolls with an enzyme cocktail prepared in RPMI 1640 media, including enzymes G and D, and reagent A from the FFPE Tissue Dissociation kit (Catalogue No. 130-118-052, Miltenyi Biotec), and an additional enzyme, LiberaseTM with a final concentration of 1 mg/mL (Catalogue No. 5401119001, Millipore Sigma). The enzyme cocktail mix was incubated at 37℃ for 15 minutes before adding to the tissue scrolls. Dissociation of each sample was carried out in a gentleMACS^TM^ Octodissociator (Miltenyi Biotec) using the “*FFPE_37C”*. The resulting cell suspension was filtered through 70-micron and 30-micron filters, centrifuged at 500g for 10 minutes at 4℃, washed with cold 1X PBS, centrifuged again, and finally resuspended in 100 μL of diluted 1X quenching buffer (provided in Chromium Next GEM Single Cell Fixed RNA Sample Preparation Kit, 10X Genomics).

Cell quality and quantity were assessed by dual-fluorescent staining with ViaStain™ Acridine Orange Propidium Iodide (AOPI) Staining Solution (Nexcelom Bioscience, Cat.No. CS2-0106) and counting on a Cellometer-K2 Automated Fluorescent Cell Counter (Nexcelom Bioscience) in Brightfield, as well as using the PE/Texas Red® channel with a bandpass filter 610/10 (fixed cells-only stain fluorescent red). The cells from each sample had minimal debris and were of high yield.

#### Probe hybridization, multiplexing, and post-hybridization washing

Two million cells from each of the four samples were individually hybridized with Human WTA Probes BC001, BC002, BC003, and BC004, (Chromium Fixed RNA Kit, Human Transcriptome, 10X Genomics), respectively, for 23 hours at 42℃ in a thermocycler. Following hybridization, samples were diluted in the post-Hyb wash buffer. Cell counts were performed using AOPI staining on the Cellometer-K2 Automated Fluorescent Cell Counter. A total of 265,000 cells from each of the four hybridized samples were then pooled equally. The pooled cells were washed three times with a post-Hyb wash buffer (initial wash with 1 mL, followed by two washes with 500 μL), each wash involving a 15-minute incubation at 42℃ and centrifugation at 850g for 5 minutes at RT. The final pellet was resuspended in 500 μL of post-Hyb resuspension buffer. All steps were performed as instructed in the Chromium Fixed RNA Profiling for Multiplexed Samples User Guide (CG000527-Rev E). Cell counting and assessment were performed using AOPI staining on the Cellometer-K2 Fluorescent Cell Counter, as previously described.

#### Single-cell GEMs formation, library construction, and sequencing

Approximately 65,000 multiplexed cells were immediately loaded with the GEM Master Mix and Gel Beads for Gel Bead-in-Emulsions (GEMs) formation, recovery, and pre-amplification, according to the manufacturer’s user guide. Flex-gene expression (Flex-GEX) libraries were constructed from 20 µL of the pre-amplified product for each sample using dual index TS-Set A by following the Chromium Fixed RNA Profiling for Multiplexed Samples User Guide (CG000527-Rev E). Library traces were analyzed on the Bioanalyzer 2100 (Agilent). The normalized Flex-GEX libraries were pooled in an equimolar ratio and sequenced on one Illumina NovaSeq 6000 SP-100 flow cell at the Molecular Biology Core Facility (MBCF), DFCI. The run parameters were 28, 10, 10, 90.

### Preprocessing and quality control

For the scRNA^1^ and snFlex datasets, barcode processing, read alignment, and UMI counting were performed using the 10x Genomics Cell Ranger pipeline (v7.2.0). Reads were aligned to the human genome reference GRCh38 (hg38) using the GENCODE v32 / Ensembl 98 annotation. snFlex probe sequences were aligned to the Flex_human_probe_v1.0.1 reference. For the snMultiome^12^ datasets, barcode processing, alignment, and UMI counting were performed using Cell Ranger ARC (v2.0.2). RNA reads were processed using Cell Ranger v7.2.0, consistent with the scRNA and snFlex datasets, and aligned to the same GRCh38 reference (GENCODE v32 / Ensembl 98). Confidently mapped, non-PCR-duplicate reads were retained to generate a gene-by-barcode count matrix for each sample.

To remove ambient RNA contamination, barcodes corresponding to low RNA content droplets were filtered from each sample using CellBender remove-background (v0.2.0)^26^. The “full” model was run with raw count matrices as input. Parameters including learning rate, number of epochs, and total droplets were optimized per sample based on the training loss curve. The expected cell count was set using Cell Ranger’s estimated cell number, rounded up to the nearest hundred. All other parameters were kept at default settings.

To identify and exclude multiplets, doublet detection was performed on the CellBender-cleaned matrices using Python Scrublet package (v0.2.2)^27^. For each sample, the expected doublet rate was set to 0.1, and doublet score thresholds were manually chosen to separate singlets from doublets. Predicted doublets were removed, resulting in singlet-only, ambient RNA-corrected count matrices for downstream analysis in Seurat (v5.1.0)^28^.

Samples corresponding to each sequencing method were merged into a single Seurat object, resulting in three separate objects representing snFlex, scRNA, and snMultiome datasets. Each object was then subjected to quality control. Genes detected in fewer than three cells and cells with fewer than 200 detected features were removed. For snFlex and scRNA datasets, cells with 5% or more of counts derived from mitochondrially encoded transcripts were excluded. Because scRNA captures whole cells, which naturally contain more cytoplasmic and mitochondrial transcripts, a more permissive threshold was applied, removing cells with 25% or more mitochondrial counts.

### Malignant cell identification and cross-cohort integration for cell type annotation

For each Seurat object, raw counts were normalized using the default log-normalization method. The top 2,000 variable genes were identified using mean-variance standardized feature values (“vst” method). Data were scaled, and principal component analysis (PCA) was performed using the top 30 principal components (PCs). To integrate samples across cohorts, we applied Harmony (v1.2.0)^29^ on the top 30 PCs, using sample identity as a covariate. The 30 Harmony-corrected dimensions were used for unsupervised Louvain clustering and visualization by uniform manifold approximation and projection (UMAP).

Malignant cells were identified based on inferred copy number variation (CNV) profiles and cluster-level expression of canonical ccRCC marker genes, such as *CRYAB, CA9, VEGFA*, and *PAX8*. For each sample, cells were subsetted from the integrated Seurat object and re-normalized and re-scaled as described above. PCA was performed on the scaled data, and the top 30 PCs were used for Louvain clustering and UMAP. Clusters were annotated as broad cell types based on the expression of canonical markers for immune, stromal, and tumor compartments. To infer CNV profiles, we used InferCNV (https://github.com/broadinstitute/inferCNV) for each sample, designating putative tumor and stromal clusters as observation groups and immune clusters as reference groups. Malignant clusters were defined based on clear patterns of copy number gain or loss, including known ccRCC features such as chromosome 3p loss. Cells within putative tumor clusters that were not labeled as malignant by InferCNV, as well as cells labeled as malignant but clustering with stromal or immune populations, were considered ambiguous and excluded from downstream analysis.

After malignant cell identification and exclusion of ambiguous cells within each sample, the resulting high-quality cells were used for downstream cross-cohort integration and cell type annotation. All samples (n = 16) were merged into a single Seurat object. The merged dataset was re-normalized, re-scaled, and subjected to PCA on the top 30 PCs. Batch correction was performed using Harmony with sample identity as a covariate. The 30 Harmony-corrected dimensions were used for Louvain clustering and UMAP visualization as described above. Broad cell types were annotated based on canonical marker expression, resulting in a final dataset of 123,357 cells spanning nine major cell types: tumor, T, myeloid, B and plasma, fibroblast, endothelial, muscle, and epithelial cells.

### Shared broad cell type identification, differential gene expression, gene set enrichment, and gene overlap analysis in the integrated object

Broad cell types shared across sequencing methods were identified from the integrated object by verifying the presence of each annotated cell type across all methods using cell type proportions and UMAP visualization. Tumor, T, myeloid, B and plasma, fibroblast, and endothelial populations were consistently detected across all methods and defined as shared cell types. The integrated object was subsetted to these shared populations, re-normalized and re-scaled following feature selection, and subjected to PCA, Harmony integration, Louvain clustering, and UMAP visualization as described above. The resulting object was annotated using canonical marker genes representing immune, stromal, and tumor compartments, yielding 122,877 high-quality cells across six shared broad cell types (tumor, T, myeloid, B and plasma, fibroblast, and endothelial).

To compare transcriptional profiles of shared cell types across methods, the integrated object containing shared cell types was further subsetted by the sequencing method. For each method, we transferred cell type annotations from the integrated object to ensure consistent labeling across datasets. We then identified cell-type specific differentially expressed genes (DEGs) for each method by performing differential expression (DE) analysis comparing cells of each Louvain cluster to all other clusters using logistic regression, including sample name as a covariate (Bonferroni-adjusted p < 0.05) (Tables S2C-E).

To evaluate whether shared markers captured comparable biological functions across methods, we defined genes identified as markers in all methods as shared markers (Tables S2B). Gene set enrichment analysis (GSEA) was performed for each cell type marker set using the clusterProfiler R package (v4.10.1)^30^. Enrichment analysis was based on gene ontology (GO) Biological Process terms, and p-values were adjusted using Benjamini-Hochberg false discovery rate (FDR) correction with a threshold of 0.05. Gene fold enrichment was calculated as the ratio of the gene ratio to the background ratio (Tables S2F).

To quantitatively assess similarities and differences in marker genes across shared cell types, pairwise gene overlap analyses were performed for each cell type using Fisher’s exact test and Jaccard similarity. Specifically, for each cell type, marker lists from each pairwise comparison (snFlex vs. scRNA, snMultiome vs. scRNA, and snFlex vs. snMultiome) were compared against an effective gene universe defined as the intersection of genes detected in both datasets. The *newGeneOverlap* and *testGeneOverlap* functions from the GeneOverlap R package (v1.38.0)^31^ were used to construct contingency tables and compute (i) the Jaccard index (proportional overlap), (ii) odds ratio (enrichment of overlap), and (iii) Fisher’s exact test p-values for statistical significance. Results were summarized per cell type with values rounded to two decimal places.

### Granular immune and tumor cell annotation in the integrated object

Annotation of granular T cell, myeloid, and tumor subtypes was performed through iterative rounds of subsetting, normalization, feature selection, scaling, PCA, Harmony integration, Louvain clustering, and DE analysis within each cell type.

For CD8⁺ T cell subtype annotation, we first subsetted the integrated object to cells belonging to the broad T cell clusters (31,162 cells). These T cells were further clustered, resulting in 10 distinct Louvain clusters. To identify CD8⁺ populations, we evaluated *CD8A* expression by calculating both the average expression level and the percentage of *CD8A*-expressing cells within each cluster. Clusters were assigned as CD8^+^ T cells if 30% or more of cells expressed *CD8A* and the average *CD8A* expression exceeded 5. A subsequent round of subsetting and re-clustering of these CD8^+^ T cells yielded six distinct CD8⁺ T cell states (14,119 cells). DE analysis was then performed by comparing each Louvain cluster to all others using a two-sided Wilcoxon rank-sum test (Bonferroni-adjusted p < 0.05). Based on cluster-specific marker genes, we identified the following CD8⁺ T cell states: 4-1BB-high, interferon-stimulated gene (ISG)-high, cycling, early activated-like, scRNA-dominant, and NK-like (Table S4A).

For tumor-associated macrophages (TAMs) a similar analytical strategy was applied. The integrated object was subsetted to the myeloid compartment and re-clustered, yielding 13 distinct Louvain subtypes (18,755 cells). For each cluster, we computed the average expression and percentage of cells expressing the TAM-associated genes *CD14, CD68*, and *CD163*. We then calculated the mean of these metrics across the three genes, defining TAM clusters as those with mean percentage expression greater than 35% and mean average expression greater than 5. These TAM clusters were further subsetted and re-clustered, identifying six TAM phenotypes (15,847 cells). DE analysis revealed the following TAM states: CXCL10-high, lipid-associated, M2-like, cycling, lncRNA-high, and lipid-associated SPP1-high (Table S4B).

For tumor subtypes, we subsetted cells annotated as tumor in the integrated object and performed re-clustering, initially identifying seven Louvain clusters (54,845 cells). Three of these were sample-specific and were excluded to ensure cross-method consistency. Re-clustering of the remaining cells identified five distinct tumor phenotypes (47,193 cells). DE analysis revealed proximal tubule-like, oxidative metabolism-high, interferon-stimulated, cycling, and lncRNA-high tumor cell states (Table S4C).

### Broad shared cell type annotation and proportional downsampling in patient 876

After identifying true malignant and high-quality cells within each method-specific object, data from patient 876 were subsetted, resulting in three Seurat objects: snFlex (7,530 cells), scRNA (10,759 cells), and snMultiome (3,004 cells). For each object, normalization, feature selection, scaling, PCA, and Louvain clustering were performed on the top 30 principal components (PCs) as previously described. Broad cell types were identified through DE analysis by comparing each Louvain cluster to all others using a two-sided Wilcoxon rank-sum test (Bonferroni-adjusted p < 0.05).

Broad cell types shared across sequencing methods were identified by visualizing the UMAP embedding of each object, in addition to calculating the proportions of each cell type. Tumor, T, myeloid, NK cell, and endothelial populations were consistently detected across all methods and defined as shared cell types. Each object was subsetted to these shared populations and re-clustered as described above. Canonical marker genes representing immune, stromal, and tumor compartments were used to re-annotate these five shared cell types, resulting in 7,138 cells in snFlex, 10,715 cells in scRNA, and 2,972 cells in snMultiome.

To enable a direct, method-level comparison within patient 876 while accounting for large differences in total cell numbers across methods, we applied a proportional downsampling strategy to snFlex and scRNA to match the total number of cells in snMultiome (2,972) while preserving their original cell type proportions. First, we calculated the proportion of each shared cell type relative to the total cell number in snFlex and scRNA. Based on the snMultiome sample size and cell type proportion, we then determined the number of cells to retain for each cell type to maintain proportional representation. A custom function (random_select_cells) was implemented to randomly select the predefined number of unique cell_id for each cell type without replacement. The selected cell identifiers were combined to generate proportionally downsampled Seurat objects for snFlex and scRNA. These downsampled datasets provided balanced representation across cell types for subsequent marker gene analysis.

### Marker gene comparison analysis in patient 876

To compare transcriptional profiles of shared cell types across methods in patient 876, we followed the same analytical framework as used for the integrated object. For each method, snMultiome, downsampled snFlex, and downsampled scRNA, we identified cell type-specific DEGs using one-versus-all logistic regression, including sequencing depth as a covariate (Bonferroni-adjusted p < 0.05) (Tables S1E-G).

To evaluate whether shared marker genes captured comparable biological functions across methods, we identified shared marker genes for each cell type and performed GSEA using the clusterProfiler R package with the same parameters as described for the integrated analysis. Enrichment analysis was based on GO Biological Process terms, with p-values adjusted using Benjamini-Hochberg FDR correction (cutoff = 0.05) as previously described (Tables S1D, H).

Finally, we assessed similarities and differences in marker gene composition across shared cell types by performing pairwise gene overlap analyses for each cell type using Fisher’s exact test and Jaccard similarity, following the same parameters and strategy described above.

### Granular immune and tumor cells analysis in patient 876

Annotation of granular T cell, myeloid, and tumor subtypes was performed through iterative rounds of subsetting, normalization, feature selection, scaling, PCA, Louvain clustering, and differential expression analysis within each cell type for each original method (non-downsampled), as described above.

For CD8⁺ T cell subtype annotation, each original object was subsetted to cells belonging to the broad T cell clusters. Due to limited cell numbers in the snMultiome dataset, detailed subclustering was performed only for snFlex and scRNA. These T cells were re-clustered, and *CD8A* expression was evaluated by calculating both the average expression level and the percentage of *CD8A*-expressing cells within each cluster. Clusters were defined as CD8⁺ T cells if 30% or more of cells expressed CD8A and the average CD8A expression exceeded 5. A subsequent round of subsetting and re-clustering of these CD8⁺ T cells identified three distinct CD8⁺ T cell states in both snFlex (687 cells) and scRNA (1,138 cells). Differential expression analysis was performed by comparing each Louvain cluster to all others using Multinomial-based Alignment of Single-cell Transcriptomics (MAST) model and two-sided Wilcoxon rank-sum tests (Bonferroni-adjusted p < 0.05). Based on cluster-specific marker genes, we identified three CD8⁺ T cell states: 4-1BB-high, 4-1BB-low, and cycling (Tables S3A-C).

To further characterize the functional states of 4-1BB-high and 4-1BB-low populations in each method, we first identified DEG by comparing 4-1BB-high and 4-1BB-low clusters using a two-sided Wilcoxon rank-sum test, setting the log fold-change threshold to 0 and the minimum expression percentage to 0.01 in each method (Tables S3D, E). Genes were then ranked by log fold-change within each method, and GSEA was performed on these ranked gene lists using the fgsea R package (v1.28.0)^32^ with the gene signature from Oliveira et al^13^.

For TAM, as described, we subsetted to the myeloid compartment in each method individually, then re-clustered the cells, and assessed the average expression and percentage of cells expressing the TAM-associated genes *CD14, CD68,* and *CD163*. The mean of these metrics across the three genes was calculated, and clusters with mean percentage expression greater than 25% and mean average expression greater than 5 were defined as TAMs. Low-quality cells were excluded. These TAM clusters were further subsetted and re-clustered, identifying three TAM phenotypes per method. DE analysis revealed lipid-associated, M2-like, and CXCR4-high TAMs in snFlex (861 cells) and scRNA (1,019 cells), and lipid-associated, CXCR4-high, and HIF1A-high TAMs in snMultiome (558 cells) (Tables 5S).

For tumor subtypes, cells annotated as tumor were subsetted from each object and re-clustered, identifying three distinct tumor clusters per method: 4,493 cells in snFlex, 4,871 cells in scRNA, and 1,621 cells in snMultiome. DE analysis was performed as described (Tables S6A-C). GSEA was then conducted on each tumor cluster marker gene set across methods using the clusterProfiler R package with the same parameters as described above. Enrichment was based on GO Biological Process terms, and p-values were adjusted using the Benjamini-Hochberg FDR correction with a threshold of 0.05. Fold enrichment was calculated as previously described (Tables S6D-F). Significant enriched pathways were then ranked by fold enrichment, and biologically relevant pathways, such as metabolic, hypoxia, renal function, and cell cycle programs, were manually selected to characterize each tumor cluster across methods.

