## Supplemental figures for "Evaluation of single-cell RNA profiling technologies using FFPE, fresh, and frozen ccRCC tumor specimens"

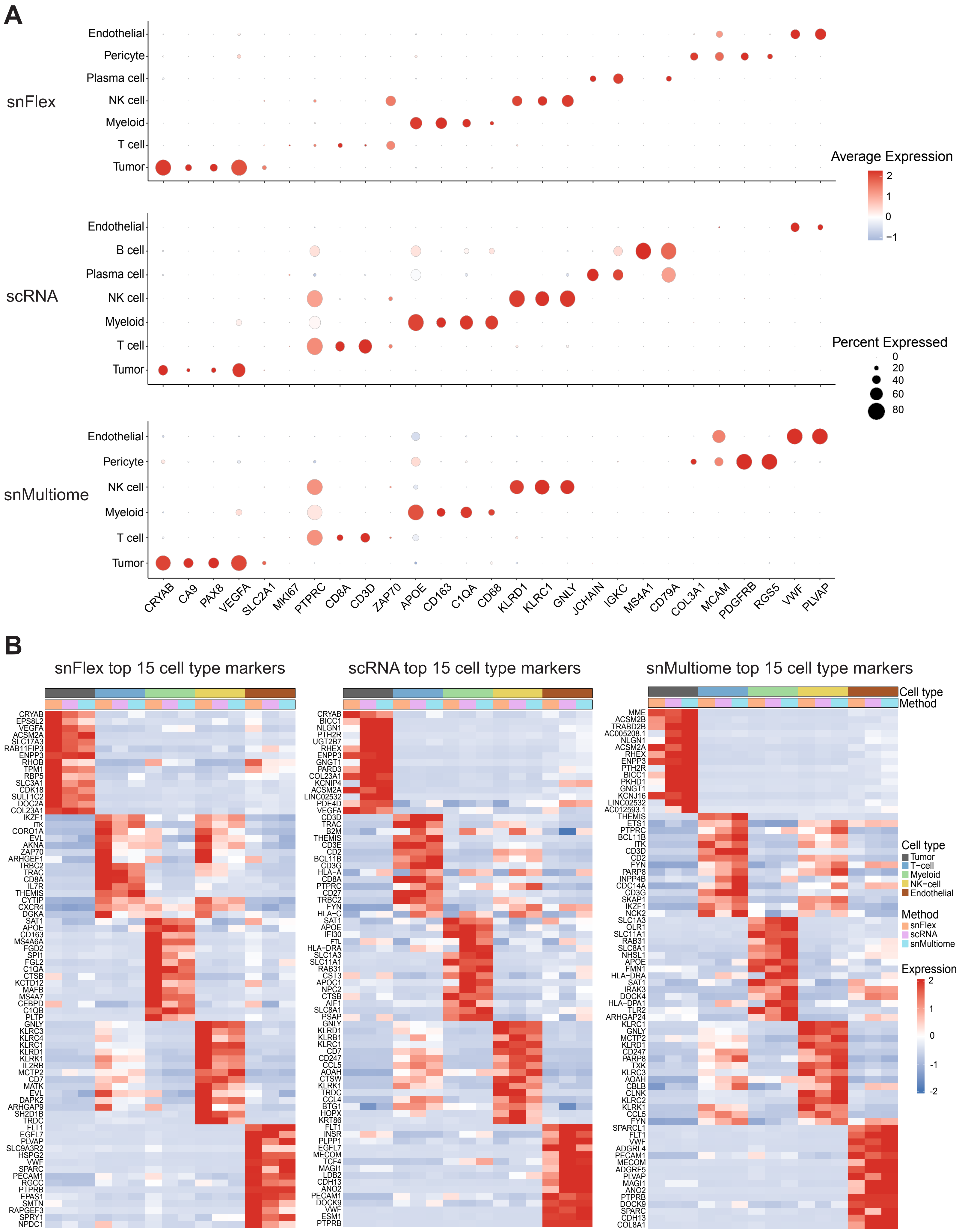
A
snFlex
Endothelial
Pericyte
Plasma cell
NK cell
Myeloid
T cell
Tumor
scRNA
Endothelial
B cell
Plasma cell
NK cell
Myeloid
T cell
Tumor
snMultiome
Endothelial
Pericyte
NK cell
Myeloid
T cell
Tumor
Average Expression
2
1
0
−1
Percent Expressed
0
20
40
60
80
CRYAB CA9 PAX8 VEGFA SLC2A1 MKI67 PTPRC CD8A CD3D ZAP70 APOE CD163 C1QA CD68 KLRD1 KLRC1 GNLY JCHAIN IGKC MS4A1 CD79A COL3A1 MCAM PDGFRB RGS5 VWF PLVAP
B
snFlex top 15 cell type markers
CRYAB EPS8L2 VEGFA ACSM2A SLC17A3 RAB11FIP3 ENPP3 RHOB TPM1 RBP5 SLC3A1 CDK18 SULT1C2 DOC2A COL23A1 IKZF1 ITK CORO1A EVL AKNA ZAP70 ARHGEF1 TRBC2 TRAC CD8A IL7R THEMIS CYTIP CXCR4 DGKA SAT1 APOE CD163 MS4A6A FGD2 SPI1 FGL2 C1QA CTSB KCTD12 MAFB MS4A7 CEBPD C1QB PLTP GNLY KLRC3 KLRC4 KLRC1 KLRD1 KLRK1 IL2RB CCL5 CD7 MATK EVL DAPK2 ARHGAP9 SH2D1B TRDC FLT1 EGFL7 PLVAP SLC9A3R2 HSPG2 VWF SPARC PECAM1 RGCC PTPRB EPAS1 SMTN RAPGEF3 SPRY1 NPDC1
scRNA top 15 cell type markers
CRYAB BICC1 NLGN1 PTH2R UGT2B7 RHEX ENPP3 GNGT1 PARD3 COL23A1 KCNIP4 ACSM2A LINC02532 PDE4D VEGFA CD3D TRAC THEMIS CD3E CD2 BCL11B CD3G HLA−A CD8A PTPRC CD27 TRBC2 FYN HLA−C SAT1 APOE IFI30 FTL HLA−DRA SLC1A3 SLC11A1 RAB31 CST3 APOC1 NPC2 CTSB AIF1 SLC8A1 PSAP GNLY KLRD1 KLRB1 KLRC1 CD7 CD247 CCL5 AOAH CTSW KLRK1 TRDC CCL4 BTG1 HOPX KRT86 FLT1 INSR PLPP1 EGFL7 MECOM TCF4 MAGI1 LDB2 CDH13 ANO2 PECAM1 DOCK9 VWF ESM1 PTPRB
snMultiome top 15 cell type markers
MME ACSM2B TRABD2B AC005208.1 NLGN1 ACSM2A RHEX ENPP3 PTH2R BICC1 PKHD1 GNGT1 KCNJ16 LINC02532 AC012593.1 THEMIS ETS1 PTPRC BCL11B ITK CD3D CD2 FYN PARP8 INPP4B CDC14A CD3G SKAP1 IKZF1 NCK2 SLC1A3 OLR1 SLC11A1 RAB31 SLC8A1 NHSL1 APOE FMN1 HLA−DRA SAT1 IRAK3 DOCK4 HLA−DPA1 TLR2 ARHGAP24 KLRC1 GNLY MCTP2 KLRD1 CD247 PARP8 TXK KLRC3 AOAH CBLB CLNK KLRC2 KLRK1 CCL5 FYN SPARCL1 FLT1 VWF ADGRL4 PECAM1 MECOM ADGRF5 PLVAP MAGI1 ANO2 PTPRB DOCK9 SPARC CDH13 COL8A1
Cell type
Method
Cell type
Tumor
T−cell
Myeloid
NK−cell
Endothelial
Method
snFlex
scRNA
snMultiome
Expression
2
1
0
-1
-2

**Figure S1. Overview of cell type and markers recovery across methods in patient 876.**

A. Dot plot showing expression of canonical cell type marker genes across cell types detected by each sequencing method. Dot size represents the percentage of cells expressing each gene within a cluster, and color indicates average expression level. The same set of marker genes is shown along the x-axis for all methods, with color scales kept consistent across panels.
B. Heatmaps showing average gene expression of the top 15 cell type-specific marker genes identified in each method for patient 876. Marker genes were determined by one-versus-all logistic regression differential expression tests (Bonferroni-adjusted $p < 0.05$), accounting for sequencing depth. For each cell type within each method, markers were ranked by adjusted p-value and log fold change, and the top 15 were selected. Average gene expression was computed on a merged Seurat object containing all methods, and each heatmap displays the top markers from one method plotted across the merged dataset. Columns are grouped by cell type and annotated by both cell type and sequencing method.

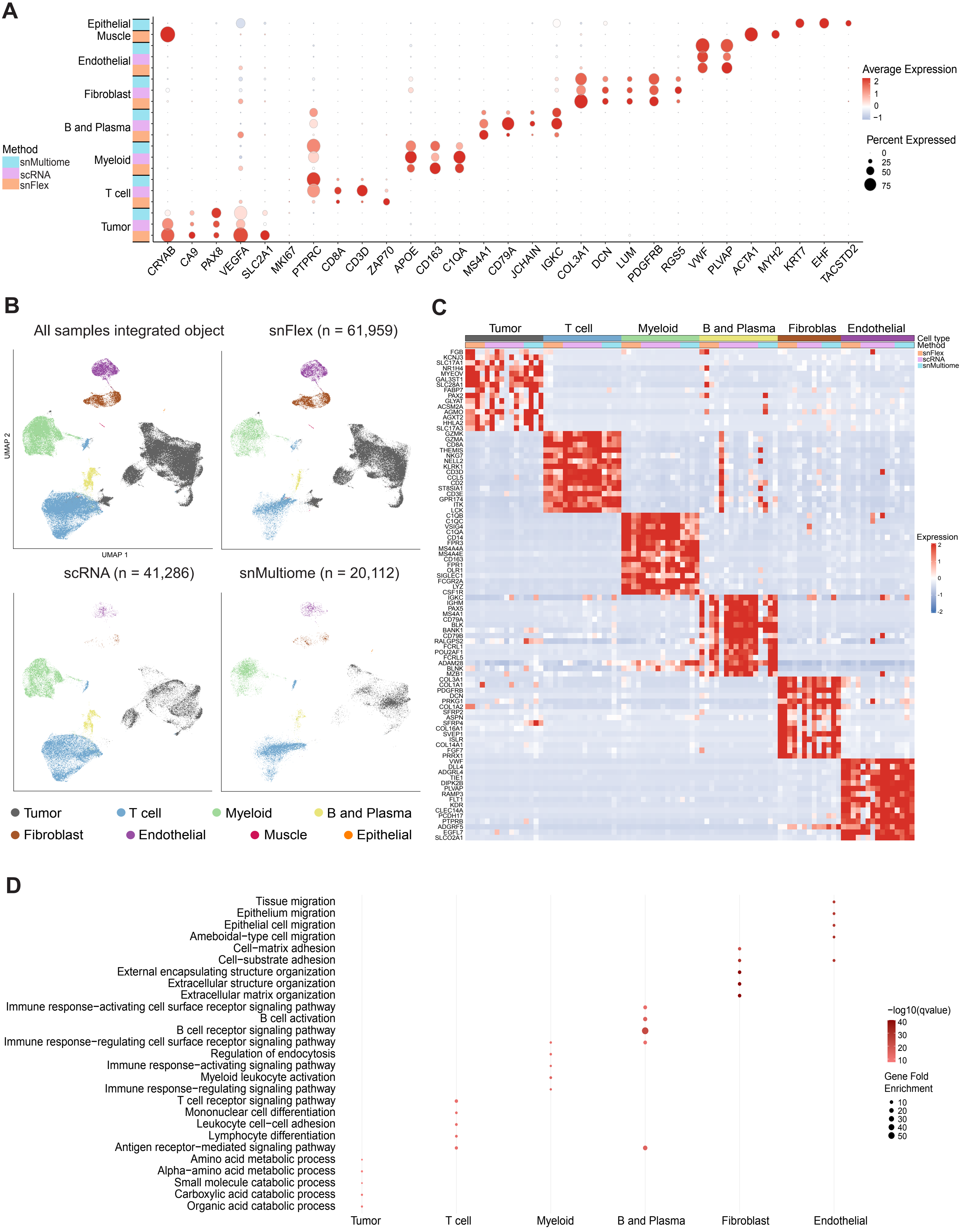
A
Epithelial
Muscle
Endothelial
Fibroblast
B and Plasma
Myeloid
T cell
Tumor
Method
snMultiome
scRNA
snFlex
CRYAB
CA9
PAX8
VEGFA
SLC2A1
MKI67
PTPRC
CD8A
CD3D
ZAP70
APOE
CD163
C1QA
MS4A1
CD79A
JCHAIN
IGKC
COL3A1
DCN
LUM
PDGFRB
RGS5
VWF
PLVAP
ACTA1
MYH2
KRT7
EHF
TACSTD2
Average Expression
2
1
0
−1
Percent Expressed
0
25
50
75
B
All samples integrated object
snFlex (n = 61,959)
scRNA (n = 41,286)
snMultiome (n = 20,112)
UMAP 2
UMAP 1
Tumor
T cell
Myeloid
B and Plasma
Fibroblast
Endothelial
Muscle
Epithelial
C
Tumor
T cell
Myeloid
B and Plasma
Fibroblas
Endothelial
Cell type
Method
snFlex
scRNA
snMultiome
Expression
2
1
0
-1
-2
D
Tissue migration
Epithelium migration
Epithelial cell migration
Ameboidal−type cell migration
Cell−matrix adhesion
Cell−substrate adhesion
External encapsulating structure organization
Extracellular structure organization
Extracellular matrix organization
Immune response−activating cell surface receptor signaling pathway
B cell activation
B cell receptor signaling pathway
Immune response−regulating cell surface receptor signaling pathway
Regulation of endocytosis
Immune response−activating signaling pathway
Myeloid leukocyte activation
Immune response−regulating signaling pathway
T cell receptor signaling pathway
Mononuclear cell differentiation
Leukocyte cell−cell adhesion
Lymphocyte differentiation
Antigen receptor−mediated signaling pathway
Amino acid metabolic process
Alpha−amino acid metabolic process
Small molecule catabolic process
Carboxylic acid catabolic process
Organic acid catabolic process
Tumor
T cell
Myeloid
B and Plasma
Fibroblast
Endothelial
−log10(qvalue)
40
30
20
10
Gene Fold
Enrichment
10
20
30
40
50

**Figure S2. snFlex, scRNA, and snMultiome capture broadly similar cell type-specific transcriptional profiles across major cell types; the same trend was detected in the full cohort analyses.**

A. Dot plot showing expression of canonical cell type marker genes across cell types from an integrated object combining all samples and sequencing methods. Cell types are grouped by method along the y-axis, with methods indicated by the color bar on the left. Dot size represents the percentage of cells expressing each gene within a cluster, and color denotes average expression level.
B. UMAP visualizations showing tumor, stromal, and immune cells from the integrated object (top left) and from each sequencing method (snFlex, scRNA, and snMultiome). Cells are colored by broad cell type, and the total number of cells detected per method is indicated in each panel.
C. Heatmap showing average gene expression of the top 15 shared marker genes for each cell type across sequencing methods, combined across all samples. Only cell types shared among methods were retained, and consistent cell type annotations from the integrated object were applied to each method. DE for each cell type was performed separately within each method using one-versus-all logistic regression tests (Bonferroni-adjusted $p < 0.05$), including sample name as a covariate. Genes identified as markers in all methods were defined as shared markers, ranked by adjusted p-value and log fold change, and the top 15 were plotted. Columns are grouped by cell type and annotated by both cell type and sequencing method.
D. Dot plot of pathways enriched among shared cell type marker genes across sequencing methods and all samples, identified with clusterProfiler based on GO Biological Process terms (Benjamini–Hochberg FDR < 0.05). Dot size indicates gene fold enrichment (GeneFoldEnrich), dot color represents statistical significance (-log10 q value). The top five enriched pathways for each cell type are displayed.

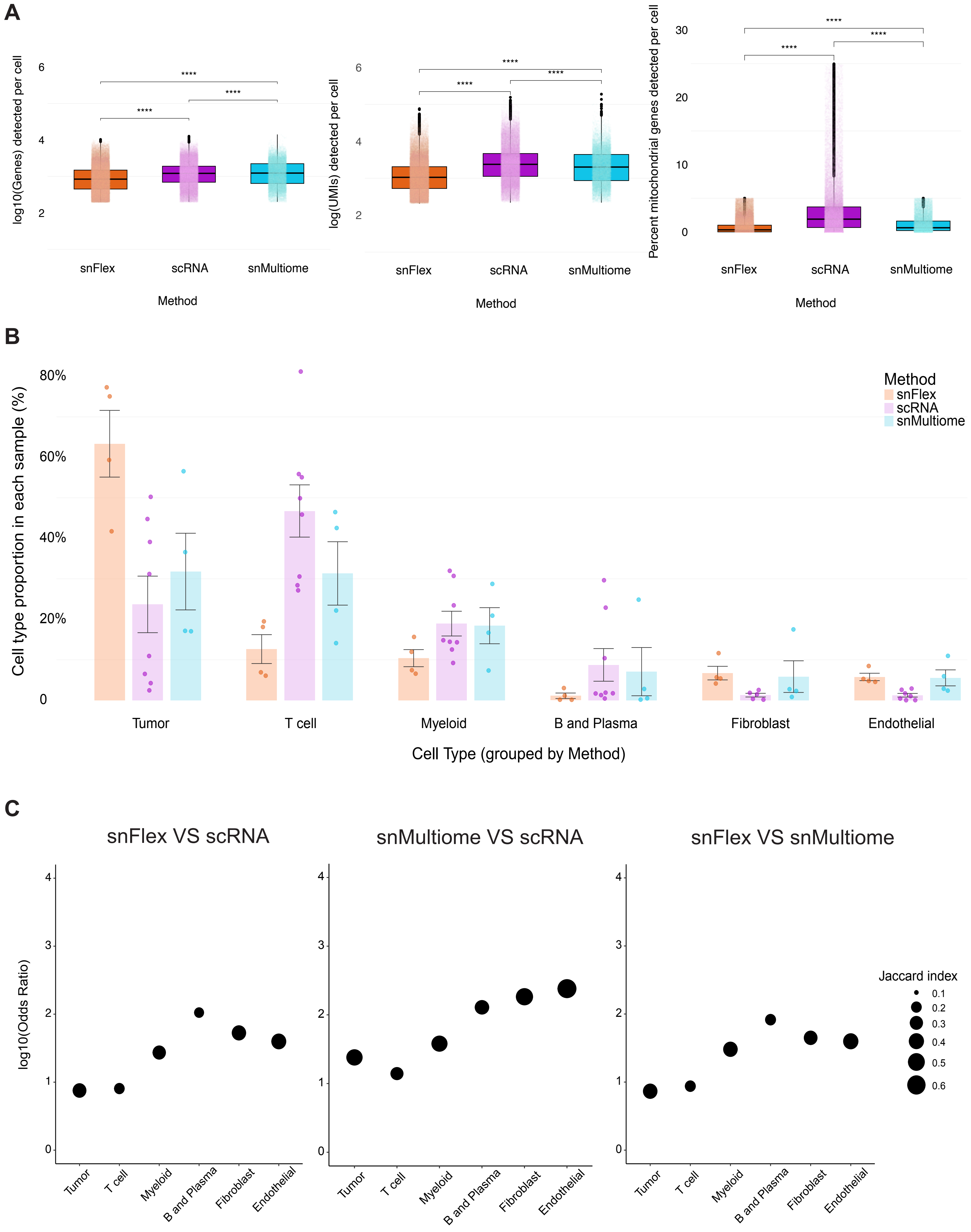
A
log10(Genes) detected per cell
6
4
2
****
****
****
snFlex
scRNA
snMultiome
Method
log(UMIs) detected per cell
6
4
2
****
****
****
snFlex
scRNA
snMultiome
Method
Percent mitochondrial genes detected per cell
30
20
10
0
****
****
****
snFlex
scRNA
snMultiome
Method
B
Cell type proportion in each sample (%)
80%
60%
40%
20%
0
Method
snFlex
scRNA
snMultiome
Tumor
T cell
Myeloid
B and Plasma
Fibroblast
Endothelial
Cell Type (grouped by Method)
C
snFlex VS scRNA
snMultiome VS scRNA
snFlex VS snMultiome
log10(Odds Ratio)
4
3
2
1
0
Tumor
T cell
Myeloid
B and Plasma
Fibroblast
Endothelial
Jaccard index
0.1
0.2
0.3
0.4
0.5
0.6

**Figure S3. Library complexity, cell type compositions, and marker expression vary across methods; the same trend was detected in the full cohort analyses.**

A. Box plots showing the number of detected genes, unique molecular identifiers (UMIs), and the percentage of mitochondrial reads per cell across sequencing methods in all samples. Gene and UMI counts are displayed on a $\log_{10}$ scale. Statistical significance was assessed using a two-sided Wilcoxon rank-sum test.

B. Bar plot showing the proportion of each cell type identified across sequencing methods in all samples. Proportions were calculated within each method and grouped by cell type; each dot represents a sample. No statistically significant differences in cell type proportions were detected using a two-sided Wilcoxon rank-sum and Kruskal-Wallis test.

C. Dot plots showing gene overlap analyses of cell type markers across sequencing methods and all samples, evaluated using Fisher's exact test and Jaccard similarity. The y-axis shows $\log_{10}$-transformed odds ratios from Fisher's exact test, representing enrichment strength. All overlaps were statistically significant ($p < 0.0001$). Dot size indicates the Jaccard index, and all panels are plotted on consistent scales.

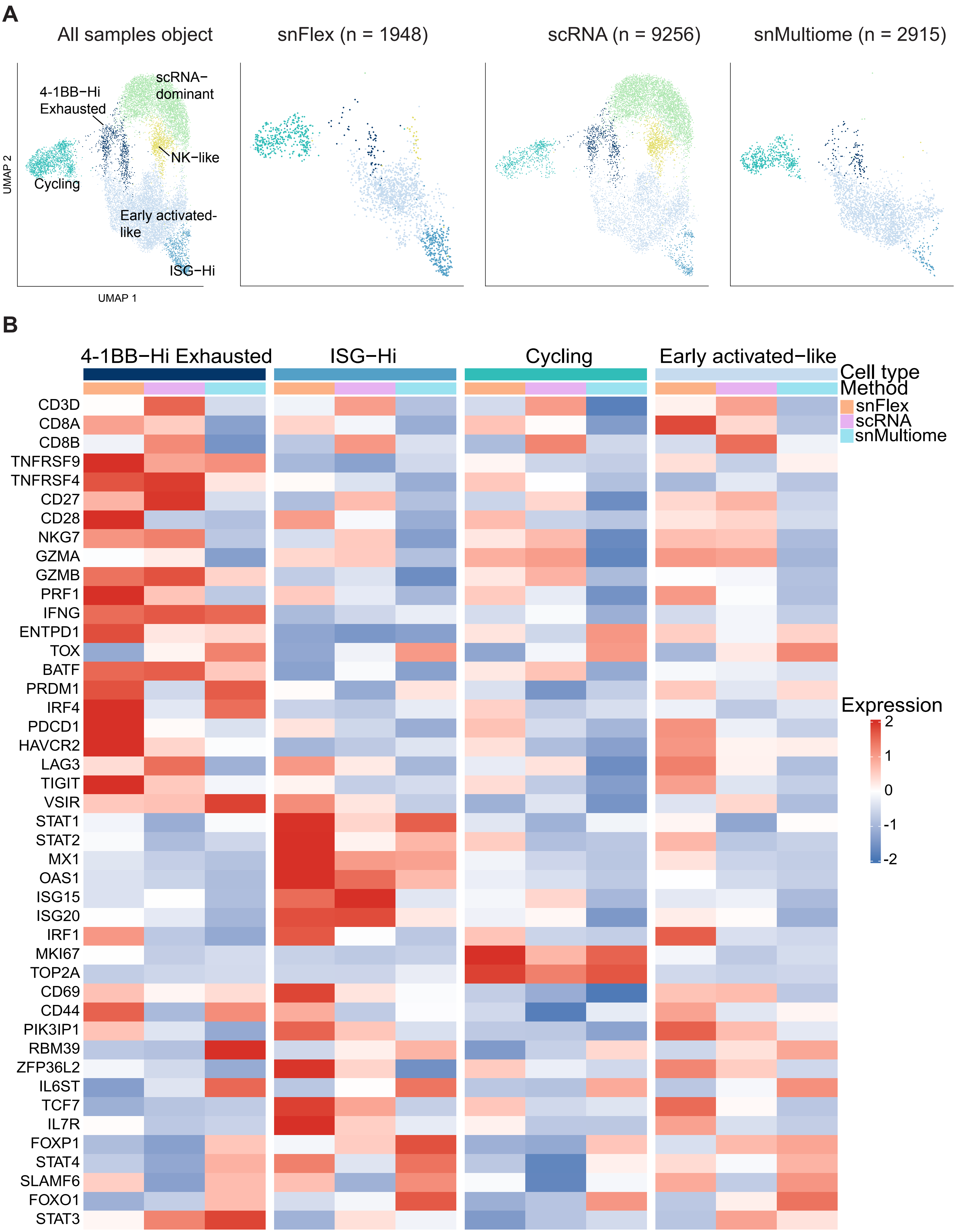
A
All samples object
snFlex (n = 1948)
scRNA (n = 9256)
snMultiome (n = 2915)
4-1BB−Hi Exhausted
scRNA−dominant
NK−like
Cycling
Early activated-like
ISG−Hi
UMAP 2
UMAP 1
B
4-1BB−Hi Exhausted
ISG−Hi
Cycling
Early activated−like
Cell type
Method
snFlex
scRNA
snMultiome
Expression
2
1
0
-1
-2
CD3D
CD8A
CD8B
TNFRSF9
TNFRSF4
CD27
CD28
NKG7
GZMA
GZMB
PRF1
IFNG
ENTPD1
TOX
BATF
PRDM1
IRF4
PDCD1
HAVCR2
LAG3
TIGIT
VSIR
STAT1
STAT2
MX1
OAS1
ISG15
ISG20
IRF1
MKI67
TOP2A
CD69
CD44
PIK3IP1
RBM39
ZFP36L2
IL6ST
TCF7
IL7R
FOXP1
STAT4
SLAMF6
FOXO1
STAT3

**Figure S4. Most CD8$^+$ T cell states are represented across the three methods.**

A. UMAP visualizations of granular CD8$^+$ T cell states from the integrated object (left) and from each sequencing method (snFlex, scRNA, and snMultiome). The total number of cells detected per method is indicated in each panel.
B. Heatmap showing scaled expression of genes defining shared CD8$^+$ T cell states identified in all methods. Marker genes were identified by a two-sided Wilcoxon rank-sum test (Bonferroni-adjusted $p < 0.05$).

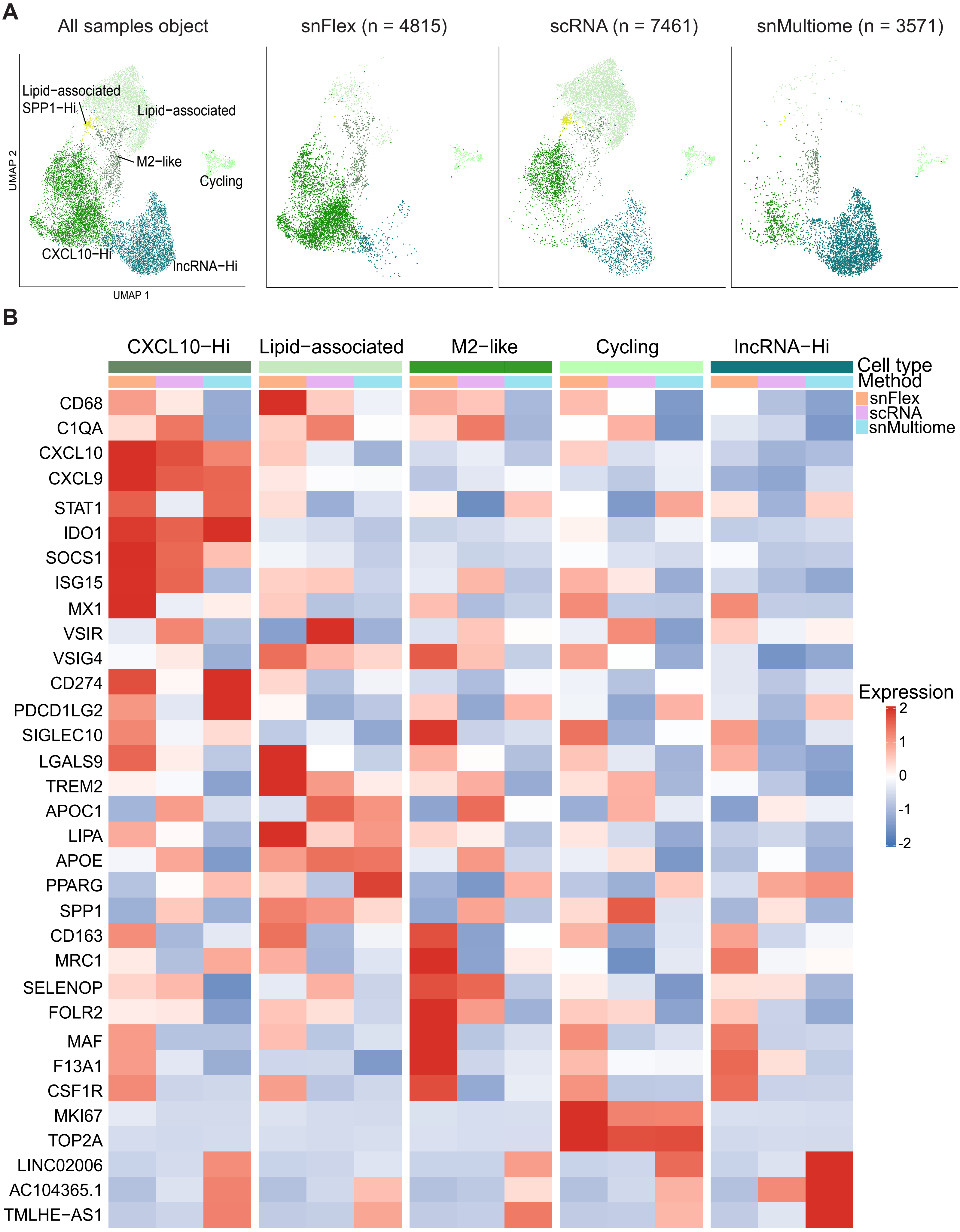
A
All samples object
snFlex (n = 4815)
scRNA (n = 7461)
snMultiome (n = 3571)
Lipid−associated SPP1−Hi
Lipid−associated
M2−like
Cycling
CXCL10−Hi
lncRNA−Hi
UMAP 2
UMAP 1
B
CXCL10−Hi
Lipid−associated
M2−like
Cycling
lncRNA−Hi
Cell type
Method
snFlex
scRNA
snMultiome
Expression
2
1
0
-1
-2
CD68
C1QA
CXCL10
CXCL9
STAT1
IDO1
SOCS1
ISG15
MX1
VSIR
VSIG4
CD274
PDCD1LG2
SIGLEC10
LGALS9
TREM2
APOC1
LIPA
APOE
PPARG
SPP1
CD163
MRC1
SELENOP
FOLR2
MAF
F13A1
CSF1R
MKI67
TOP2A
LINC02006
AC104365.1
TMLHE−AS1

**Figure S5. TAMs states are broadly concordant across methods.**

A. UMAP visualizations of granular TAM cell states from the integrated object (left) and from each sequencing method (snFlex, scRNA, and snMultiome). The total number of cells detected per method is indicated in each panel.

B. Heatmaps showing scaled expression of cell type markers defining shared TAM granular cell types identified across methods in all samples, identified by two-sided Wilcoxon rank-sum test (Bonferroni-adjusted $p < 0.05$).

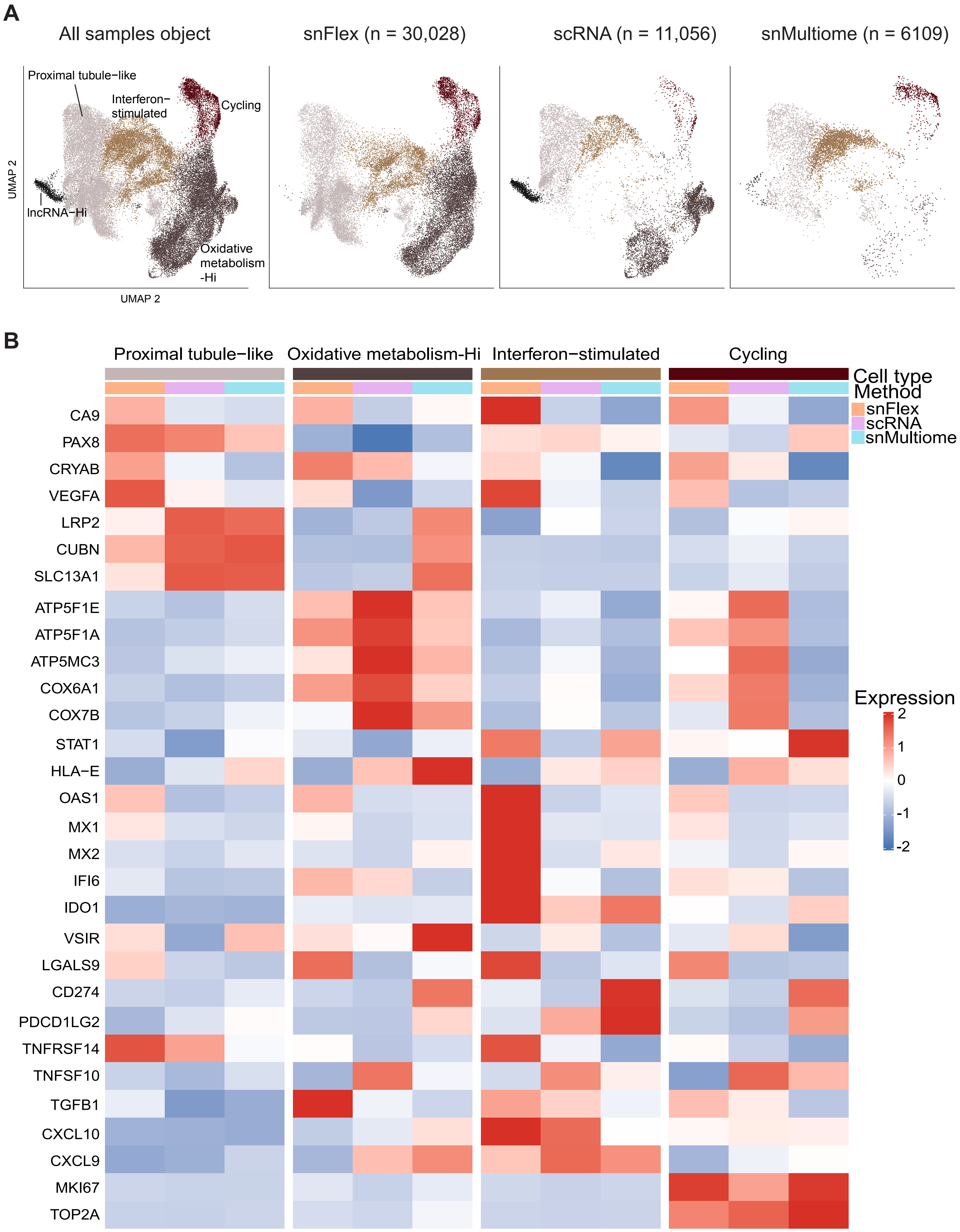
A
All samples object
snFlex (n = 30,028)
scRNA (n = 11,056)
snMultiome (n = 6109)
Proximal tubule−like
Interferon−stimulated
Cycling
lncRNA−Hi
Oxidative metabolism -Hi
UMAP 2
UMAP 2
B
Proximal tubule−like
Oxidative metabolism−Hi
Interferon−stimulated
Cycling
Cell type
Method
snFlex
scRNA
snMultiome
Expression
2
1
0
-1
-2
CA9
PAX8
CRYAB
VEGFA
LRP2
CUBN
SLC13A1
ATP5F1E
ATP5F1A
ATP5MC3
COX6A1
COX7B
STAT1
HLA−E
OAS1
MX1
MX2
IFI6
IDO1
VSIR
LGALS9
CD274
PDCD1LG2
TNFRSF14
TNFSF10
TGFB1
CXCL10
CXCL9
MKI67
TOP2A

**Figure S6. Each method supports a relatively similar phenotypic characterization of ccRCC tumor clusters.**

A. UMAP visualizations of granular tumor cell clusters from the integrated object (left) and from each sequencing method (snFlex, scRNA, and snMultiome). The total number of cells detected per method is indicated in each panel.
B. Heatmaps showing scaled expression of marker genes defining granular tumor programs across sequencing methods in shared tumor cell types identified in all methods. Marker genes were identified using a two-sided Wilcoxon rank-sum test (Bonferroni-adjusted $p < 0.05$).
